# *Clostridioides difficile* utilizes siderophores as an iron source and FhuDBGC contributes to ferrichrome uptake

**DOI:** 10.1101/2023.09.29.560173

**Authors:** Jessica L. Hastie, Hannah L. Carmichael, Bailey M. Werner, Kristin E. Dunbar, Paul E. Carlson

## Abstract

*Clostridioides difficile* remains a public health threat commonly observed following antibiotic use. Due to the importance of iron in many cell processes, most bacteria, including *C. difficile*, have multiple mechanisms of acquiring iron. Previous studies have examined ferrous iron uptake in *C. difficile*, here we focus on the role of siderophores. In a growth assay, we show that *C. difficile* can use a variety of siderophores as the sole iron source. In *C. difficile,* two ABC transporters induced under low iron conditions are predicted siderophore importers: FhuDBGC and YclNOPQ. We hypothesized that these transporters are responsible for the uptake of the siderophores we tested. To investigate the specificity of these transporters, we purified the substrate binding proteins and examined siderophore binding using thermal shift. We demonstrate increased stability between one siderophore binding protein, FhuD, and the siderophore ferrichrome, suggesting a binding interaction. This specificity correlates with the inability of an *ΔfhuDBGC* mutant to grow efficiently under iron limiting conditions in the presence of ferrichrome. While *C. difficile* used additional siderophores in our growth experiments, we did not observe increased thermal stability between the receptor proteins and any of the other siderophores tested, suggesting these siderophores do not bind these receptors and other siderophore import mechanisms remain to be elucidated. Redundancy in iron acquisition is a microbial survival adaptation to cope with the constant battle for iron within a host. Greater knowledge about how *C. difficile* acquires iron will provide insight about how *C. difficile* colonizes and persists in the colon.

**IMPORTANCE:** This study is the first example of *C. difficile* growing with siderophores as the sole iron source and describes the characterization of the ferric hydroxamate uptake ABC transporter (FhuDBGC). This transporter shows specificity to the siderophore ferrichrome. While not required for pathogenesis, this transporter highlights the redundancy in iron acquisition mechanisms which *C. difficile* uses to compete for iron during an infection.

## INTRODUCTION

*Clostridioides difficile* is most well-known for causing diarrhea after antibiotic use in both hospital- and community-acquired settings (1). This Gram-positive spore forming bacterium persists in the environment until reaching the gastrointestinal tract where germination and outgrowth occur. This outgrowth and subsequent colonization require the acquisition of sufficient nutrients, including iron. Metals are a vital resource that must be carefully managed to avoid both excess and insufficiency. Additionally, at the host-pathogen interface, multiple host metal storage mechanisms act to starve bacteria of nutrient metals in a process known as nutritional immunity (2). Due to the importance of many trace metals in bacterial cells processes, it is crucial to understand how bacterial pathogens acquire metals. These processes are often complex, with multiple mechanisms providing overlapping function to maintain intracellular metal pools at appropriate levels, while competing with host cells and other members of the microbiota for these resources.

In the gastrointestinal tract, an environment where fluctuations in oxygen and pH can impact the state of iron, the main form of iron is ferrous iron (3). While *C. difficile* encodes multiple ferrous iron transporters, the *feo1* system, encoded by Cd630_RS08100 – Cd630_RS08115, is the most highly induced in iron starved conditions (4–6). Further, deletion of *feoB1 (Cd630_RS008110* also known as *Cd630_14790)* results in decreased intracellular iron, increased metronidazole resistance, and in some strains, decreases toxin production (7). Other genes upregulated in low iron conditions include a zinc transporter, metabolic genes, and two predicted siderophore transporters. While previous work showed *C. difficile* can use many types of iron as an iron source, siderophores were not included in this study (8) and the role of siderophores in *C. difficile* iron acquisition remains unclear.

Siderophore production and/or utilization is an advantageous means of acquiring iron for both pathogenic and commensal bacteria. Siderophores are small molecules, with a very high affinity for iron (K_f_ >10^30^), that bacteria disperse to scavenge for iron (9). The diversity of siderophores is predicted to be vast (>500 types of siderophores), and siderophores are categorized based on general structure into four groups: hydroxamate, catecholate, phenolate, or mixed (9, 10). While siderophore biosynthesis genes have been identified in some Clade 3 *C. difficile* genomes, most strains of *C. difficile*, including the lab strain Cd630, do not encode for siderophore biosynthesis genes (11). The presence of two putative siderophore transporters, in the absence of siderophore biosynthesis genes, suggests most *C. difficile* strains likely use siderophores produced by other members of the gut microbiota, also known as xenosiderophores. It is unknown how xenosiderophore acquired iron impacts the ability of *C. difficile* to compete with other bacteria and host cells for iron during an infection.

Here, we show that *C. difficile* 630 is able to utilize an assortment of siderophores *in vitro*, some more efficiently than others, as sole iron sources. Of the two putative siderophore transporters identified in *C. difficile*, only the siderophore binding protein from the FhuDBGC transporter, FhuD, displayed specificity for any of the 10 siderophores tested. When incubated with ferrichrome, FhuD showed increased protein stability. Further, deletion of the *fhuDBGC* transporter showed impaired growth compared to the growth of the wild-type strain when ferrichrome was provided as a sole iron source. Iron acquisition in *C. difficile* is an important process layered with redundancy. This work highlights the ability of *C. difficile* to use siderophore as an iron source and the transporter-siderophore specificity between FhuDBGC and ferrichrome.

## MATERIALS AND METHODS

### Bacterial strains, media and growth conditions

Strains and primers used in this study are listed in Supplemental Tables 1 and 2. *C. difficile* strains were grown in an anaerobic chamber (10% hydrogen, 5% CO_2_, 85% N_2_) (Coy Lab Products, MI) at 37°C in brain heart infusion (BD Life Sciences) supplemented with 0.5% yeast extract and 0.1% cysteine (BHIS) unless otherwise stated. *E. coli* strains were grown in Luria-Bertani (LB, BD Life Sciences) at 37°C supplemented with antibiotics as indicated. Antibiotics were used at the following concentrations: for *C. difficile*, thiamphenicol (15 µg/ml); for *E. coli,* ampicillin (100 µg/ml), chloramphenicol (20 µg/ml), and kanamycin (50 µg/ml).

### Mutant construction

Unless otherwise noted, all PCR was performed using Phusion High-Fidelity DNA polymerase (ThermoScientific). Flanks for the genes of interest were synthesized by GenScript to contain 1000 bp on each side of the gene and inserted into plasmid pMSR0 (12) following digestion with PmeI resulting in plasmid pJLH123, which was transformed into HB101/pRK24 (13). Plasmids were conjugated into *C. difficile* as previously described (14) and transconjugants were selected for on BHIS plus D-cycloserine (250 µg/ml), cefoxitin (8 µg/ml), and thiamphenicol (15 µg/ml). Primary insertion of the plasmid into the chromosome was confirmed by PCR with combinations of JLHP127/617/609/610 and then streaked on BHIS plus anhydrotetracycline (100 ng/µl) to select for the secondary crossover event. Colonies were patched and screened by PCR with JLHP127/617 for deletion of the gene of interest. Whole genome sequencing confirmed deletion of the genes.

### Construction of complementation plasmids

Plasmid pJLH150 was constructed by PCR amplifying the sequence of *fhuDBCG+MATE* with the native promoter (∼200 bp), using JLHP836/838 and cloned into BamHI and NheI digested pDSW1278 (15) by Gibson assembly (16) using NEBuilder HiFi DNA Assembly Master Mix (NEB). This plasmid was conjugated into *C. difficile* as described above.

### Iron restricted growth curves

Overnight cultures of *C. difficile* were back-diluted into iron starved conditions (BHIS plus 75 μM 2’2’-dipyridyl) until the cells reached mid-log phase. Cells (5 mL) were centrifuged (Flexifuge, max speed) for 5 minutes, washed with PBS (5 mL) and centrifuged again. The resulting pellet was resuspended in 1 mL of iron depleted media (IDM, supplemental table 3) and adjusted to an OD_600_ of 0.5. These cells were used to inoculate (1:10) 96 well plates containing the media of interest (IDM + siderophore). Growth at 37°C was monitored by OD_600_ over 12 hours in a Tecan Sunrise plate reader (shaking prior to reading at hourly intervals).

### Siderophores

Siderophores were purchased from Biophore Research Products and resuspended to make a 2 mM stock concentration. The following siderophores were resuspended in water: ferrichrome, enterobactin, salmochelin, yersiniabactin, aerobactin, schizokinen, arthrobactin, and coprogen. Pyochelin was resuspended in ethanol and vibriobactin was resuspended in methanol. All siderophores were further diluted with water, buffer, or media as appropriate for the experiment.

### Construction of expression plasmids

The extracellular domain of FhuD (AA 23-313) was PCR amplified with JLHP373/374. The plasmid pET24a+ (MilliporeSigma) was PCR amplified with JLHP375/376 and digested with DpnI. The resulting PCR products were assembled using Gibson assembly resulting in plasmid pJLH55.

### Protein purification

*E. coli* BL21 cells containing plasmid pJH55 were grown overnight at 37°C in LB, back-diluted 1:20 into MagicMedia (Invitrogen), and then grown at 30°C for 24 hours. Cells were harvested by centrifugation (10,000 xg, 10 minutes, RT) and frozen at −80°C until purification. Cell pellets were thawed on ice, resuspended in 25 mL BugBuster Master Mix (MilliporeSigma), and incubated rocking for 20 minutes. Cellular debris was removed by centrifugation (10,000 xg, 10 minutes, 4°C). Strep-Tactin XT resin (1 mL) was prepared following the manufacturer’s instructions for gravity flow purification (Iba Life Sciences). After loading the cleared lysate, the column was washed with 10 column volumes of buffer W and eluted in 0.5 mL fractions with buffer BXT. The fractions were analyzed by SDS-PAGE (Any kD Mini-PROTEAN TGX gel, Bio-Rad) and quantified (Qubit Protein, Invitrogen).

### Thermal Shift

Purified FhuD (10 μM) was incubated with increasing concentrations of ferrichrome (2-200 μM), FeSO_4_ 7H_2_O (2-200 μM), or other siderophores (100 μM) and analyzed on a Prometheus NT.48 by heating from 20°C to 95°C with a ramp rate of 1°C/min. The fluorescent intensity was measured at 330 nm and 350 nm.

### Animal experiments

Mice were pretreated with cefoperozone (0.5 mg/mL) in their drinking water for five days followed by normal water for two days prior to oral gavage with 100 μl of 10^6^ *C. difficile* spores as previously described (17, 18). Mice were monitored daily for changes in weight and symptoms (hunched, scruffy fur) and fecal pellets were collected daily to plate for CFU/g on BHIS plates containing taurocholate (1%), cefoxitin (8 µg/ml), and D-cycloserine (250 µg/ml). Mice were sacrificed on day 4. The animals used in this study were on an approved protocol overseen by the Food and Drug Administration Animal Care and Use Committee.

## RESULTS

### *C. difficile* can utilize siderophore as a sole iron source

To examine siderophore dependent growth in *C. difficile*, we performed growth curves in iron limited conditions. First, we cultured the bacteria in rich media with a chelator (BHIS + 75 µM 2-2-dipyridyl) to starve the cells of iron. After washing away the media containing the chelator, we back-diluted cells into a defined iron depleted media (IDM) supplemented with 2 µM iron or 2 µM siderophore preloaded with iron. We tested an assortment of siderophores from three siderophore types: hydroxamate, catecholate, and phenolate. The siderophores arthrobactin, ferrichrome, vibriobactin, enterobactin, salmochelin, yersiniabactin and pyochelin restored the ability of *C. difficile* to grow comparable to iron supplementation (Figure 1A-C). The siderophores aerobactin, coprogen and schizokinen showed intermediate restoration of growth (Figure 1A). These results demonstrate that siderophores are an additional mechanism of iron acquisition for *C. difficile*.

**Figure 1.**
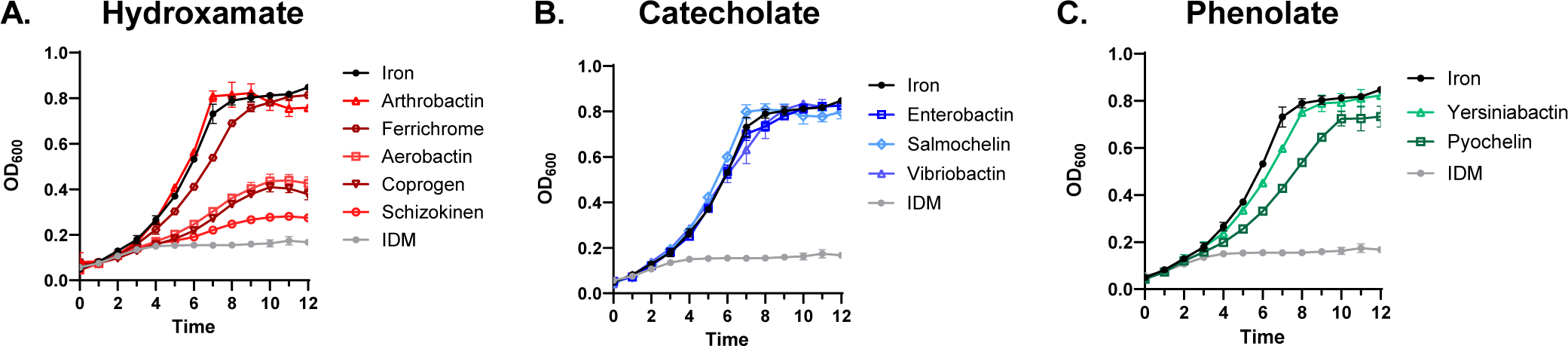
*C. difficile* utilizes multiple types of siderophore as a sole iron source in iron limiting conditions. *C. difficile* 630 was grown in iron starved conditions (BHIS + 75 μM 2’2’-dipyridyl) before inoculation into iron depleted media (IDM, gray) supplemented with 2 μM FeSO_4_ (black) or 2 µM of siderophore pre-loaded with iron (A. hydroxamate siderophores, B. catecholate siderophores, or C. phenolate siderophores). Bacterial growth was assessed by determining the OD_600_ hourly over 12 hours. Data are presented as the mean ± standard deviation of three technical replicates. Data are representative of three biological replicates.

### Characterization of putative siderophore ABC transporters

Multiple studies in *C. difficile* (4–6) have identified two ABC transporters predicted to be involved in siderophore uptake that are upregulated in iron limiting conditions. Both transporters are preceded by Fur binding sites, suggesting transcriptional control by the iron uptake regulator Fur (4). The first transporter is composed of Cd630_RS153360/*fhuD* (substrate binding protein), Cd630_RS15355/*fhuB* and Cd630_RS28769/*fhuG* (permeases), and Cd630_RS28750/*fhuC* (ATPase). A fifth gene Cd630_RS15350 (MATE family efflux transporter) is directly downstream of *fhuDBGC* (Figure 2A). To test if Cd630_RS15350 is co-transcribed with *fhuDBGC*, we performed intergenic RT-PCR and found Cd630_RS15350 is in an operon with fhuDBGC (Figure 2B). Despite being in an operon with the transporter genes, it is unclear if Cd630_RS15350 is involved in the function of the FhuDBGC transporter. The second predicted siderophore transporter is composed of Cd630_RS08975/*yclN* and Cd630_RS08980/*yclO* (permeases), Cd630_RS08985/*yclP* (ATPase), and Cd630_RS08990/*yclQ* (substrate binding protein).

**Figure 2:**
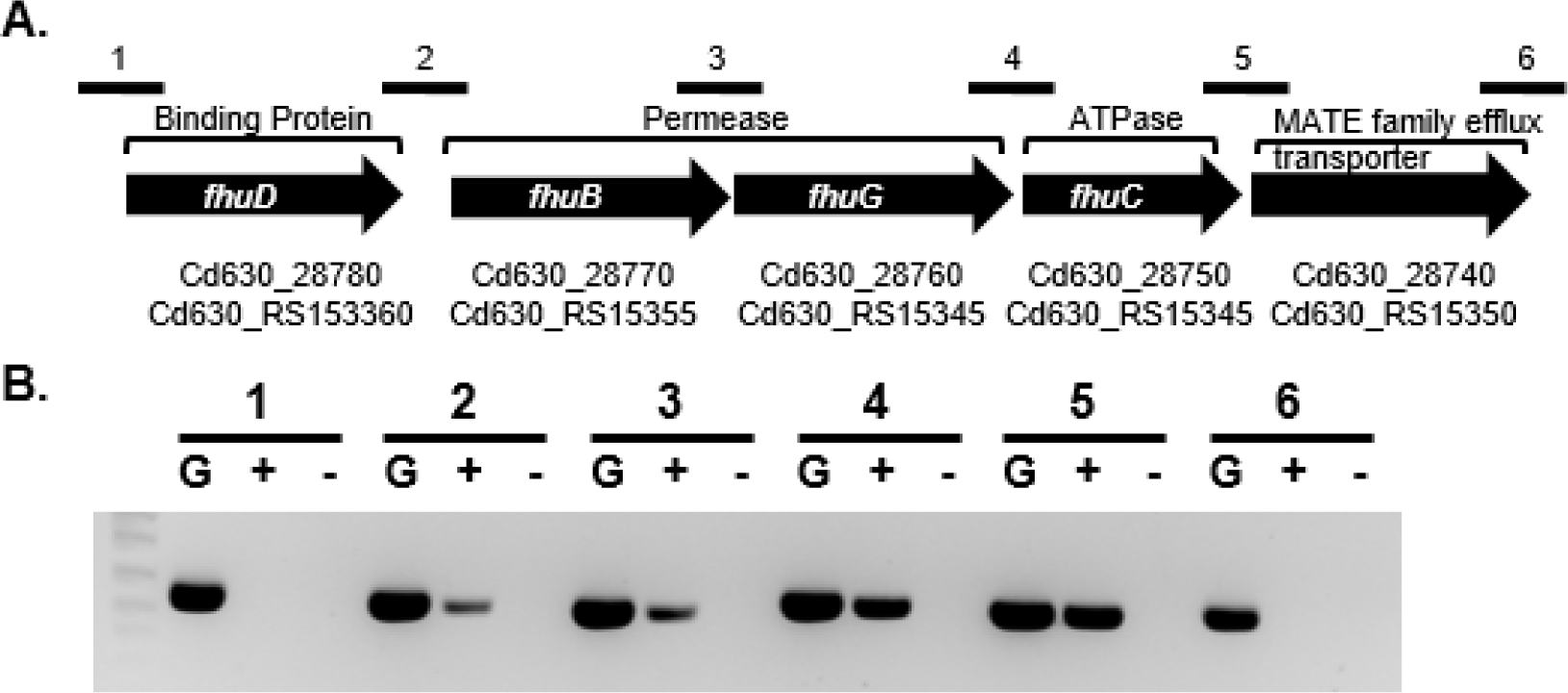
Characterization of FhuDBGC. Gene layout of transporter *fhuDBGC +MATE* (A). Reverse-Transcriptase Polymerase Chain Reaction (RT-PCR) analysis of the intergenic regions of *fhuDBGC + MATE* was performed using primer sets (1–6) described in Table S2 to detect mRNA at the regions indicated. Amplified cDNA was analyzed by electrophoresis. G = genomic DNA, (+) = RT added, (-) = no RT added (B).

Based on homology, FhuDBGC is predicted to import hydroxamate siderophores and YclNOPQ to import catecholate siderophores. To further investigate attributes of the siderophore binding proteins, we modeled their structure using AlphaFold2 version 2.1.2 (19) and aligned their sequence with homologs from other organisms using Clustal Omega version 1.2.4 (20). The predicted structures from both substrate binding proteins contain two lobes connected by a single linker, typical of class three substrate binding proteins, which is most common for siderophore binding proteins (21) (Supplemental Figure 1). Despite general conserved structural attributes, siderophore binding proteins often show specificity towards structurally similar siderophores. The Fhu transporter is conserved in several organisms; the most well-characterized examples are in *Escherichia coli, Bacillus subtilis* and *Staphylococcus aureus*. In the Gram-negative *E. coli*, several outer membrane receptors (FhuA, FhuE, and IutA) capture hydroxamate siderophores and, with the aid of TonB, move them across the outer membrane to FhuD, found in the periplasmic space (22). In *E. coli,* FhuD facilitates the transport of aerobactin, ferrichrome, and coprogen across the inner membrane (23). Alternatively, in *B. subtilis* FhuD has only been implicated in recognizing ferrichrome (24). *S. aureus* encodes two copies of FhuD: FhuD1, which only transports ferrichrome and ferrioxamine B, and FhuD2, which transports ferrichrome, ferrioxamine B, aerobactin, and coprogen (25). We aligned *C. difficile* FhuD with the corresponding proteins from these organisms using Clustal Omega. The percent identity of *C. difficile* FhuD ranged from 16.9% with *E. coli* to 26.3-28.9% with *B. subtilis* and *S. aureus* (Supplemental Figure 2). The specificity of *C. difficile* FhuD is unclear considering some hydroxamate siderophores (arthrobactin and ferrichrome) restore growth to the level of iron supplementation, while others (aerobactin, coprogen and schizokinen) only partially restore growth (Figure 1).

We compared the YclNOPQ transporter to well-known catecholate importers in *B. subtilis*, *Bacillus anthracis and E. coli*. In *B. subtilis*, the siderophore binding protein YclQ (also known as FpiA) is responsible for recognizing petrobactin (26) and is 45.2% identical to *C. difficile* YclQ (Supplemental Figure 3). *B. subtilis* is unable to synthesize petrobactin, but another Gram-positive organism *B. anthracis* synthesizes both petrobactin and encodes two siderophore binding proteins (FatB and FpuA) that bind petrobactin *in vitro* (27). However, in *B. anthracis* only FpuA is required for growth in iron limited conditions (28). The catecholate siderophores we tested (enterobactin, salmochelin and vibriobactin) are usually produced by Gram-negative bacteria. In *E. coli*, several outer membrane receptors have been identified that interact with catecholate molecules (FepA, IroN, Fiu, FecA and Cir) (*29*). Enterobactin for example, is recognized by FepA, and is then transported across the outer membrane to the substrate binding protein FepB, which binds enterobactin in the periplasm to initiate transport across the inner membrane (30). We compared *C. difficile* YclQ to these substrate binding proteins (Supplemental Figure 3), and the percent identity of YclQ to FepB, FpuA and FatB ranged from 20.0% to 26.3%. Based on these observations we predict *C. difficile* YclQ may recognize petrobactin; however, it is unclear if YclQ may also be responsible for importing other catecholate siderophores such as enterobactin, salmochelin and vibriobactin.

### ABC transporter FhuDBGC shows binding specificity to ferrichrome

We hypothesized each transporter is responsible for the uptake of specific siderophores. Previous work with FhuDBGC in other organisms suggests FhuD is responsible for the capture of ferrichrome. To test if FhuD interacts with ferrichrome, we purified the extracellular domain of FhuD using a Strep-Tactin tag and analyzed the stability of the protein by nano differential scanning fluorimetry (nanoDSF). With increasing concentrations of ferrichrome (2µM −200 µM) the melting temperature of FhuD increases, suggesting stabilization of FhuD (Figure 3A). The pH of the elution buffer used with Strep-Tactin purification is pH 8. Considering the physiological pH of the colon is 5.5-7.0 (31), we adjusted the pH of FhuD using HCl to pH 6 and 7. Decreasing the pH caused FhuD to unfold at a lower temperature (Figure 3B). However, the stabilization of FhuD with ferrichrome was consistent at all pH’s tested and the iron-free form, desferrichrome, did not increase the melting temperature of FhuD (Figure 3C). These results indicate FhuD interacts with iron-loaded ferrichrome.

**Figure 3.**
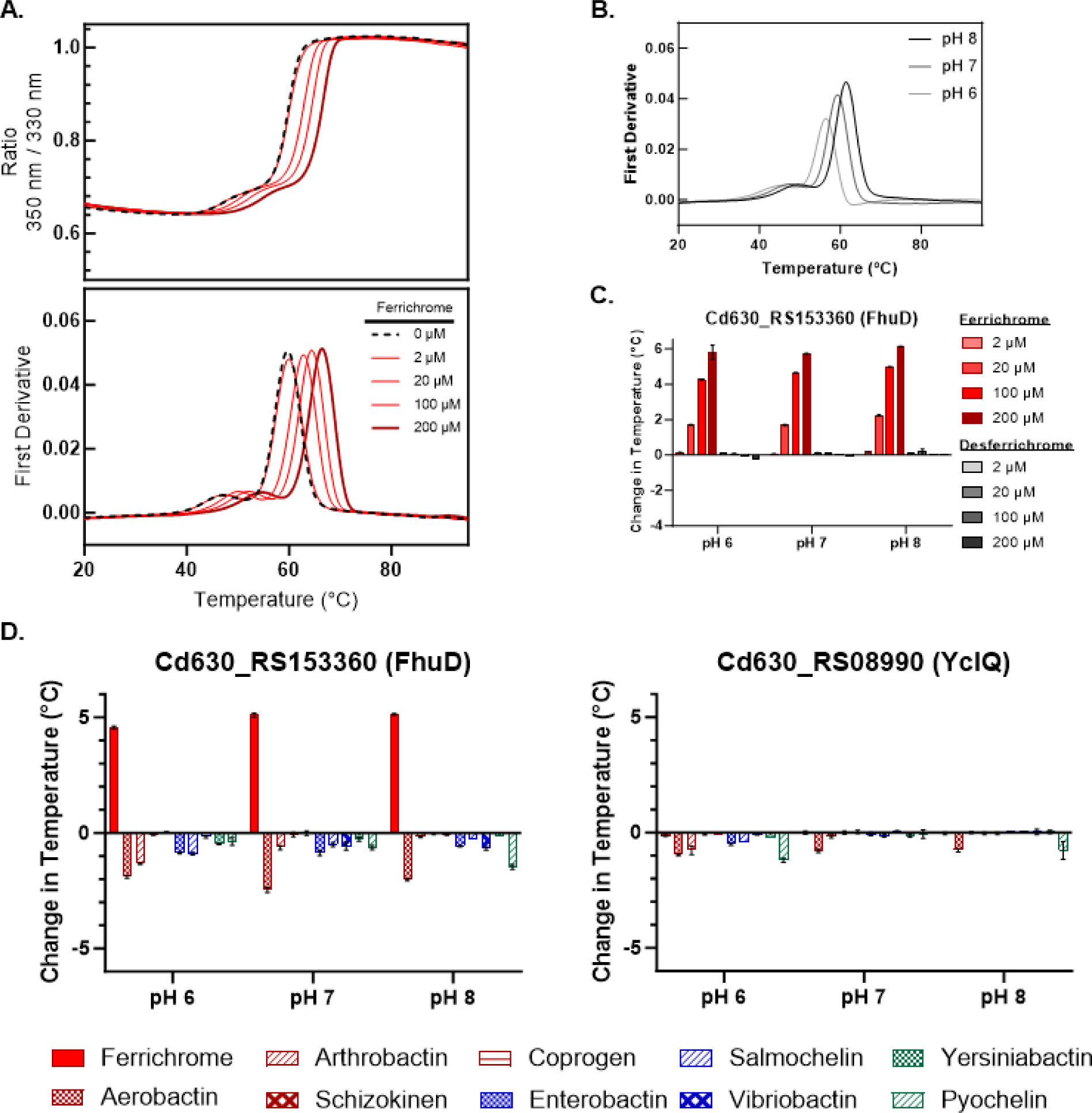
Ferrichrome binds FhuD. Thermal unfolding traces of FhuD (10 μM) with increasing concentrations of ferrichrome (2-200 μM) (A) or across pH 6-8 (B). See Supplemental Figure 4 for unfolding traces at each pH for each protein (10 μM) without siderophore used to calculate the change in unfolding temperature. FhuD with a gradient of ferrichrome and desferrichrome. Data are representative of multiple protein preparations, performed in technical triplicate, and error bars represent the standard error of the mean (C). Purified FhuD and YclQ (10 μM) with different siderophores at 100 μM. Data are representative of one protein preparation, performed in technical triplicate, and error bars represent the standard error of the mean (D).

To further examine transporter-siderophore specificity, we tested the unfolding of FhuD with other siderophores (enterobactin, salmochelin, vibriobactin and yersiniabactin) including several hydroxamates siderophores (aerobactin, arthrobactin, schizokinen, and coprogen). None of the other siderophores caused an increase in protein stability comparable to ferrichrome (Figure 3D). These results confirm that the change in FhuD protein unfolding is specific to ferrichrome, which is consistent with FhuD from *B. subtilis* which specifically recognizes ferrichrome (32).

To determine if the YclQ interacts with our panel of siderophores, we purified the extracellular domain of YclQ, similar to FhuD, using a Strep-Tactin tag. None of the siderophores tested caused YclQ stabilization, suggesting these siderophores do not interact with YclQ (Figure 3D). It is possible that YclQ, like FhuD, is highly specific for a siderophore that we have not tested, such as petrobactin.

### Deletion *of fhuDBGC* impairs ferrichrome uptake

Based on the increased stability of FhuD in the presence of ferrichrome, we hypothesized cells without the FhuDBGC transporter would be unable to utilize ferrichrome as a sole iron source. We used allelic exchange to make a clean deletion of *Cd630_RS153360-Cd630_RS15350 (*including the gene annotated as the MATE family efflux transporter, but we refer to this knockout as *ΔfhuDBGC* for simplicity). The *fhuDBGC* mutant grew comparable to WT with iron supplementation (Figure 4A) and was impaired, but not abolished, in IDM media supplemented with ferrichrome (Figure 4B). We constructed the vector pJLH150 to complement Δ*fhuDBGC* expressed under the operon’s native promoter. Plasmid complementation of *fhuDBGC* restores ferrichrome utilization comparable to WT (Figure 4B). Uptake of the other siderophores was not impacted by the deletion of *fhuDBGC* (Figure 5, Supplemental Figure 5). We note the mutant constantly shows slightly improved growth with desferrichrome, schizokinen, coprogen and pyochelin. It is possible deletion of *fhuDBGC* leads to the upregulation of other iron acquisition systems such as the Feo transporters that result in slightly enhanced growth, perhaps due to contaminating free iron in the iron-loaded siderophores. These results demonstrate the FhuDBGC transporter is necessary for optimal utilization of ferrichrome.

**Figure 4.**
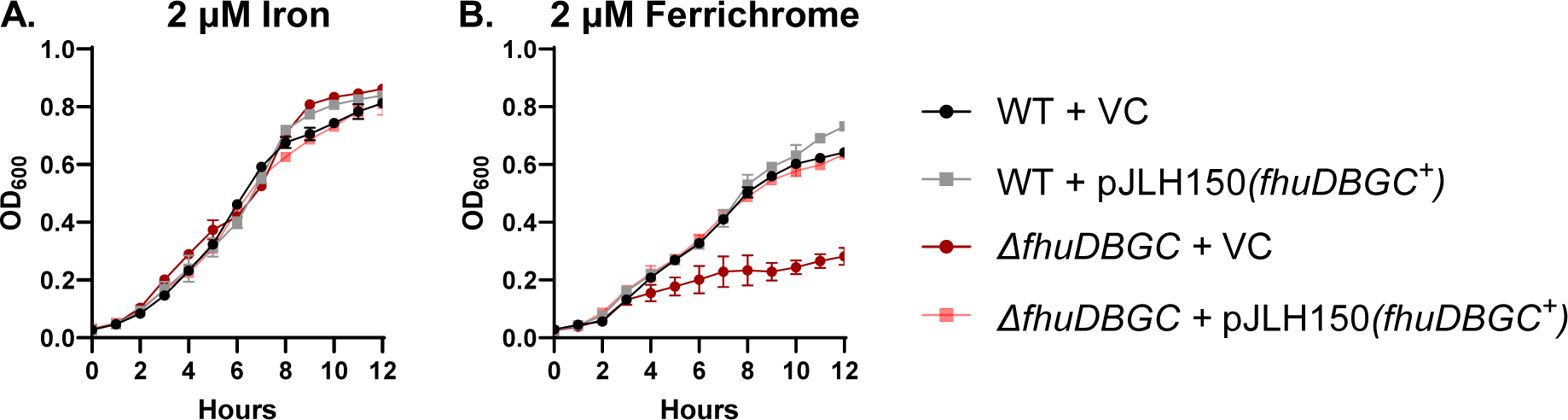
Growth of *ΔfhuDBGC* is impaired in the presence of ferrichrome. WT Cd630 and its isogenic fhuDBGC transporter mutant containing VC or pJLH150 were grown in iron starved conditions (BHIS + 75 μM 2’2’-dipyridyl). The cells were standardized to an OD of 0.5 before diluting 1:10 into iron depleted media (IDM) supplemented with 1% xylose and 2 μM FeSO_4_ (A) or 2 µM of ferrichrome pre-loaded with iron (B). Bacterial growth was assessed by determining the OD_600_ hourly for 12 hours. Data are the mean ± standard deviation of three technical replicates. Data are representative of three biological replicates. VC = vector control.

**Figure 5.**
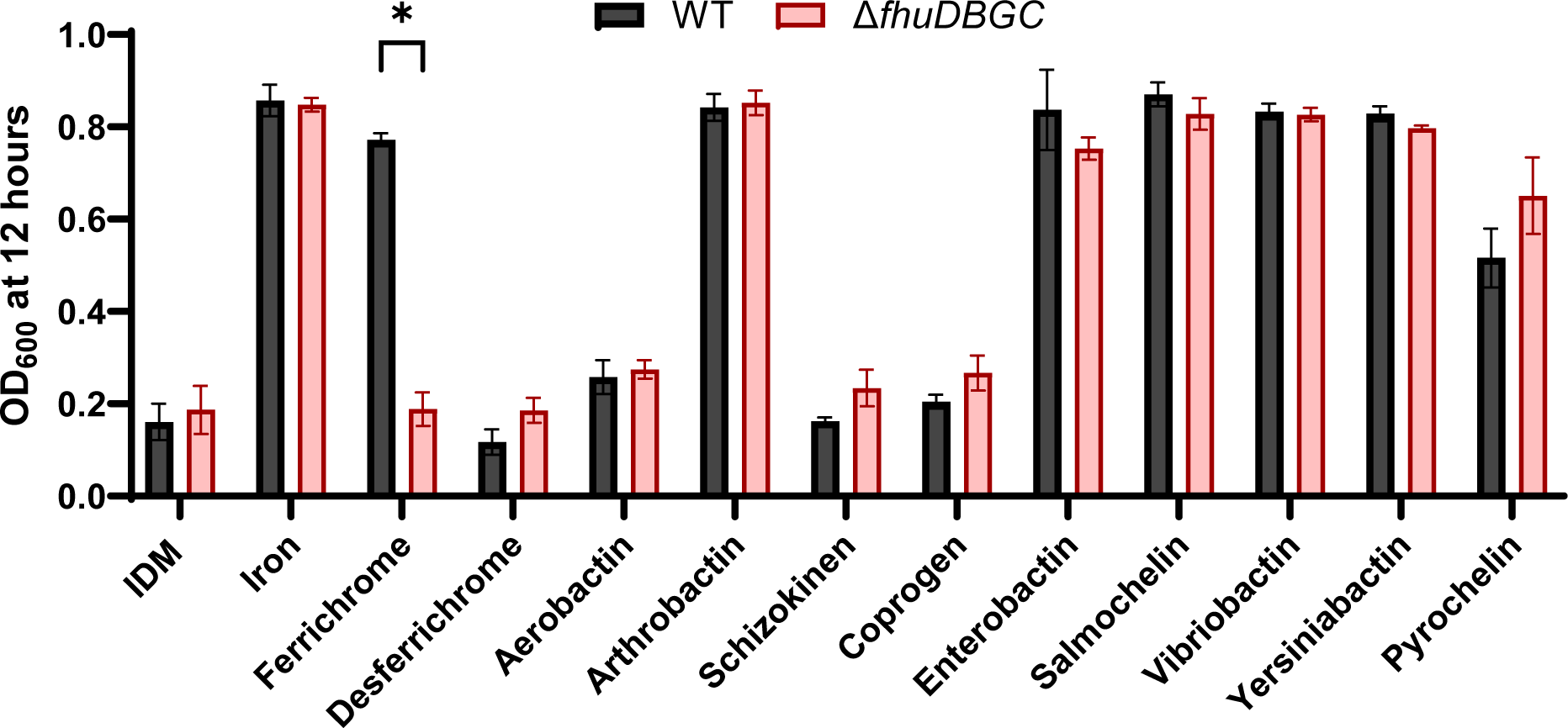
Reduced growth of *ΔfhuDBGC* is specific to ferrichrome. *C. difficile* 630 (black) and its isogenic *fhuDBGC* transporter mutant (red) was grown in iron starved conditions (BHIS + 75 μM 2’2’-dipyridyl) and adjusted to an OD 0.5 before diluting 1:10 into IDM, IDM supplemented with 2 μM FeSO_4_, or IDM supplemented with 2 µM of siderophore pre-loaded with iron. Bacterial growth was assessed by determining the OD_600_ hourly for 12 hours. Data are presented as the endpoint OD_600_ at 12 hours mean ± standard deviation of three technical replicates. Data are representative of three biological replicates. An unpaired *t* test was used to determine statistical significance (* = p <0.001).

### Deletion of *fhuDBGC* alone does not impact pathogenesis

Previous work has shown that *C. difficile fhuB* was upregulated during hamster infection (*4*) and growth under microaerophilic conditions (33) suggesting the FhuDBGC transporter may be important during colonization. To determine the impact of the *ΔfhuDBGC* mutation on colonization, we used the cefoperazone mouse model of *C. difficile* infection (CDI) (17). We infected mice with ∼10^5^ spores of wild-type Cd630 or Cd630 *ΔfhuDBGC* and monitored the mice for colonization by plating fecal samples and counting CFUs over four days. The mice infected with *ΔfhuDBGC* remained colonized comparable to WT for the duration of the experiment, suggesting removal of *fhuDBGC* alone does not impair iron acquisition enough to impact colonization (Figure 6).

**Figure 6.**
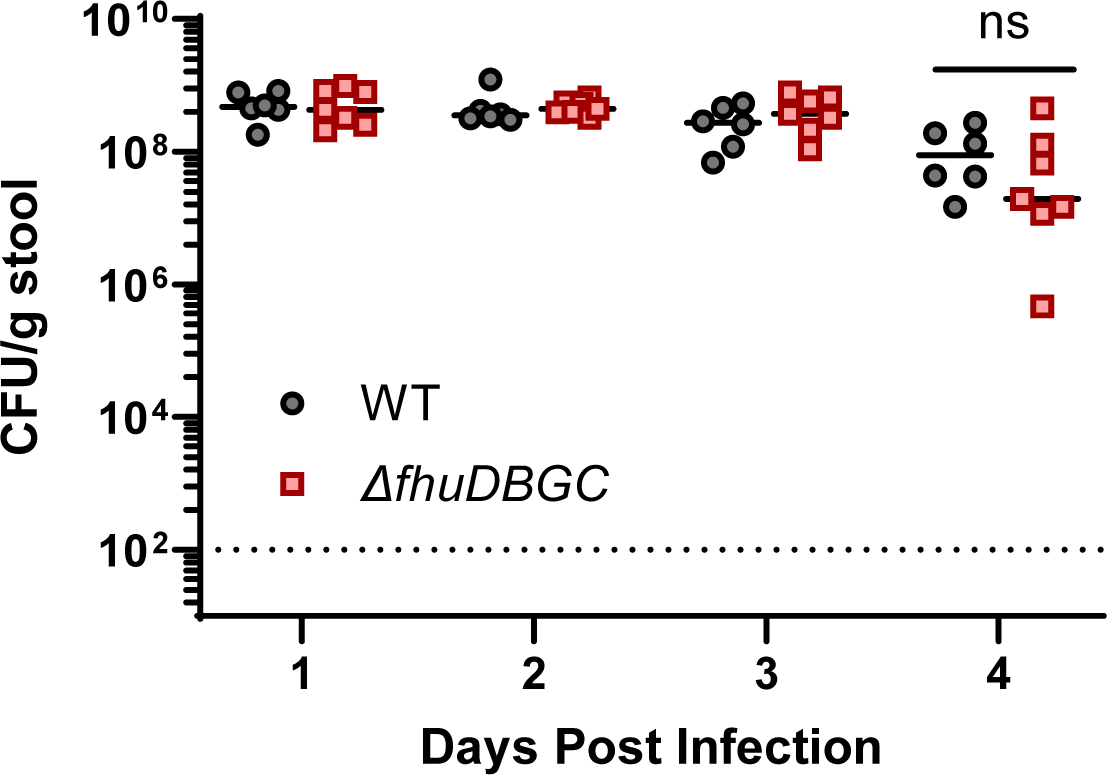
Deletion of *fhuDBGC* does not impair murine gut colonization. *C. difficile* colonization in the cefoperazone treated mouse model of CDI. Mice were pretreated with cefoperazone in the drinking water for five days, changed to normal water for two days, then inoculated with ∼ 10^5^ spores by oral gavage. *C. difficile* colonization was assessed by plating stool for four days post infection (WT infected mice n=6, Δ*fhuDBGC* infected mice n=7). An unpaired *t* test was used to determine statistical significance, where ns = not significant. The dashed line represents the limit of detection for the assay.

## DISCUSSION

Since the *C. difficile* genome encodes genes for putative siderophore transporters, which are induced under iron limiting conditions, many have speculated that this pathogen exploits xenosiderophores as an iron source. In this study, we sought to characterize *C. difficile* siderophore utilization and showed that *C. difficile* can use a variety of siderophores as an iron source. These results indicate that siderophores represent an additional iron acquisition redundancy that *C. difficile* likely uses to compete with host cells and other commensals while colonizing the colon. Using a biochemical approach, we identified siderophore-transporter specificity between FhuD and ferrichrome. This interaction was further confirmed *in vitro* using a *fhuDBGC* mutant that was unable to efficiently grow when ferrichrome was provided as the sole iron source. Although we only tested a small subset of siderophores, we did not see any interaction between YclQ and siderophores tested, suggesting additional mechanisms beyond FhuD and YclQ are involved in siderophore uptake.

The amount and availability of siderophores within the gut environment is unclear. Considering that most of the siderophores we tested restored growth of *C. difficile* comparable to iron, it appears *C. difficile* is a prolific xenosiderophore user. In addition to bacteria, plants and fungi produce siderophores. Ferrichrome, in particular, is synthesized by yeast (*Schizosaccharomyces pombe*), fungi (*Ustilago maydis*) and bacteria (*Lactobacillus casei ATCC334*)(34–36). Another gastrointestinal pathogen *Salmonella enterica* utilizes fungal siderophores, which are thought to be present in the gastrointestinal tract from both live fungi found in the microbiota and acquired through diet (37). Given the vast diversity of siderophores it is likely *C. difficile* can utilize many siderophores beyond the scope that we tested.

Although several studies have highlighted the competitive advantage siderophores provide to pathogens and commensals competing for nutrients in the gut (38, 39), few studies have examined the role of siderophore acquired iron in anaerobic organisms. Rocha et al. showed that *Bacteroides vulgatus* and *Bacteroides thetaiotaomicron* can grow using enterobactin and salmochelin as an iron source. They additionally showed *Bacteroides fragilis* uses ferrichrome independently of FhuD (40). In *Clostridium kluyveri*, Seedorf et al. identified a siderophore biosynthetic locus and further showed a color change in the Chrome Azurol S (CAS) assay when the strain was grown on iron restricted media (41). The genome of Cd630 does not encode for siderophore biosynthesis genes, but a siderophore locus has been identified in genomes of some *C. difficile* isolates (11) suggesting that, similar to aerobic bacteria, the majority of anaerobes may use xenosiderophores, while select strains produce their own siderophore(s).

Siderophores are best known as chelators with an extremely strong affinity for ferric iron. The coordinated iron molecule is thought to be released from the siderophore upon the reduction of iron from its ferric to ferrous form in the presence of a reducing environment, with the help of a reductase or cleavage from the siderophore. We cannot rule out the possibility that our observed growth restoration in the presence of siderophore is due the reduction of the siderophore bound ferric iron upon prolonged exposure in the anaerobic chamber, which would then allow uptake by the ferrous uptake systems (Feo). However, this is unlikely for the following reasons: First, just as there is great diversity in siderophore structure, some siderophores bind iron stronger than others. Enterobactin, which is thought to have the strongest interaction with ferric iron (K = 10^52^ M^-1^ (42)) restores growth well. Within the other siderophores we tested, some restored growth well, while others restored growth less efficiently, suggesting *C. difficile* has some preference regarding siderophore use. Although some studies have compared siderophore-iron affinity, none have examined all the siderophores used in this study using the same technique. It is possible some of these siderophores (aerobactin, schizokinen and coprogen) have a weaker affinity for iron compared to others we tested and are not actually taken up by *C. difficile*. Second, to address the role of oxygen (the anaerobic environment facilitating iron reduction), we pre-incubated ferrichrome for 24 hours in the anaerobic chamber. This anaerobic ferrichrome did not improve the growth of the *fhuDBGC* mutant in IDM media (data not shown) suggesting siderophore restored growth is not due to the uptake of reduced iron via the Feo transporters.

Bacteria often have several receptors to recognize different siderophores. Considering the broad siderophore utilization observed in *C. difficile*, it is unlikely FhuD and YclQ are the only mechanisms responsible for siderophore capture. Purified FhuD was highly specific for ferrichrome, and purified YclQ did not interact with any of the siderophores we tested. While pH did influence the unfolding temperature of each siderophore binding protein, it did not alter the interaction between the protein and siderophore. At all pH’s tested, only ferrichrome caused FhuD protein stabilization. We cannot rule out the possibility that protein destabilization indicates binding through the masking of tryptophan or tyrosine residues, but the majority of the siderophores (except ferrichrome) caused some destabilization; therefore we consider this possibility to be unlikely. We were unable to obtain purified petrobactin, the substrate of *B. subtilis* YclQ, but we hypothesize YclQ is responsible for petrobactin acquisition. A third transporter Cd630_RS15975-Cd630_RS15960 was observed to be upregulated in low iron conditions but is predicted to be involved in aliphatic sulfonate or nitrate/sulfonate/bicarbonate transport. The first gene in this operon (Cd630_RS15975) is predicted to be an iron hydrogenase, which is a complex metalloenzyme (43). Low iron conditions may induce this operon due to the importance of this iron hydrogenase to acquire iron or respond to limited iron pools. Future work will aim to identify the receptors for siderophores other than ferrichrome.

*C. difficile* modulates internal and surrounding metal pools using a variety of mechanisms including the Feo transporters, a heme efflux pump (HatT), a heme sequestering protein (HsmA), and transporters for other metals (ZupT – zinc, OppA – nickel) (44). Herein, we present an additional mechanism using the transporter FhuDBGC to acquire the siderophore ferrichrome. The *fhuDBGC* mutant did not cause attenuation in the CDI mouse model likely because acquisition mechanisms other than FhuDBGC were functional. Elucidating how siderophore acquired iron influences *C. difficile’s* pathogenesis and colonization will require eliminating multiple iron acquisition mechanisms. Despite these challenges, *C. difficile* exploits an assortment of siderophores, likely including many siderophores beyond those we tested, which may provide a competitive advantage during infection.

## Acknowledgements

This work was supported by the Intramural Research Program of the Center for Biologics Evaluation and Research, Food and Drug Administration. This project was supported in part by an appointment to the Research Fellowship Program at the OVRR/CBER, U.S. Food and Drug Administration, administered by the Oak Ridge Institute for Science and Education through an interagency agreement between the U.S. Department of Energy and FDA. We thank Michael Schmitt, Eric Peng and members of the Carlson lab for helpful comments on the manuscript.

## Supplementary Information

**Supplemental 1.**
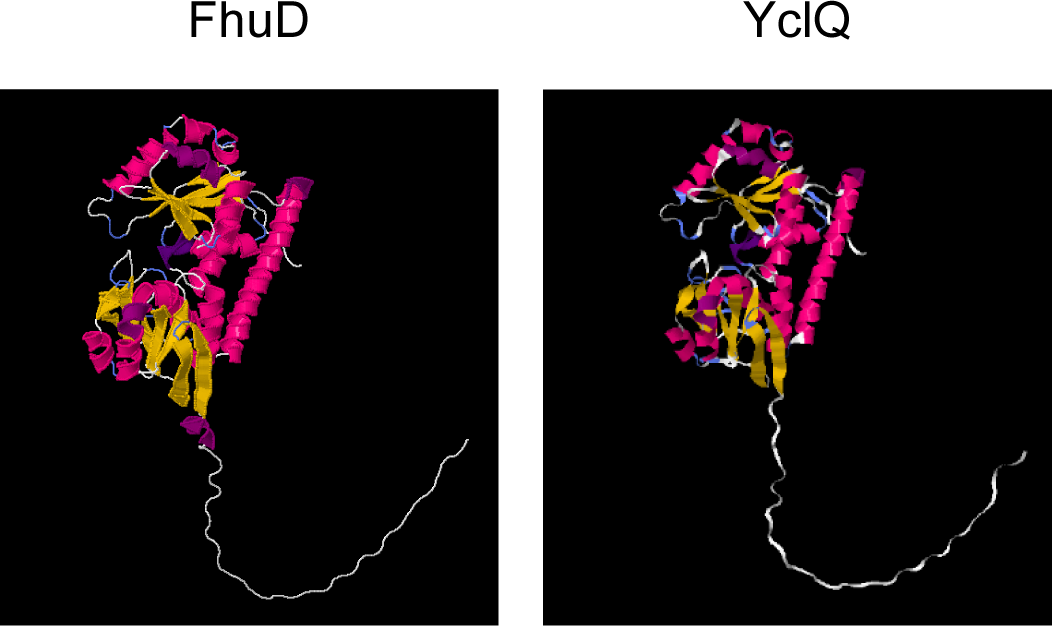
AlphaFold2 predicted structures of the siderophore binding proteins.

**Supplemental 2.**
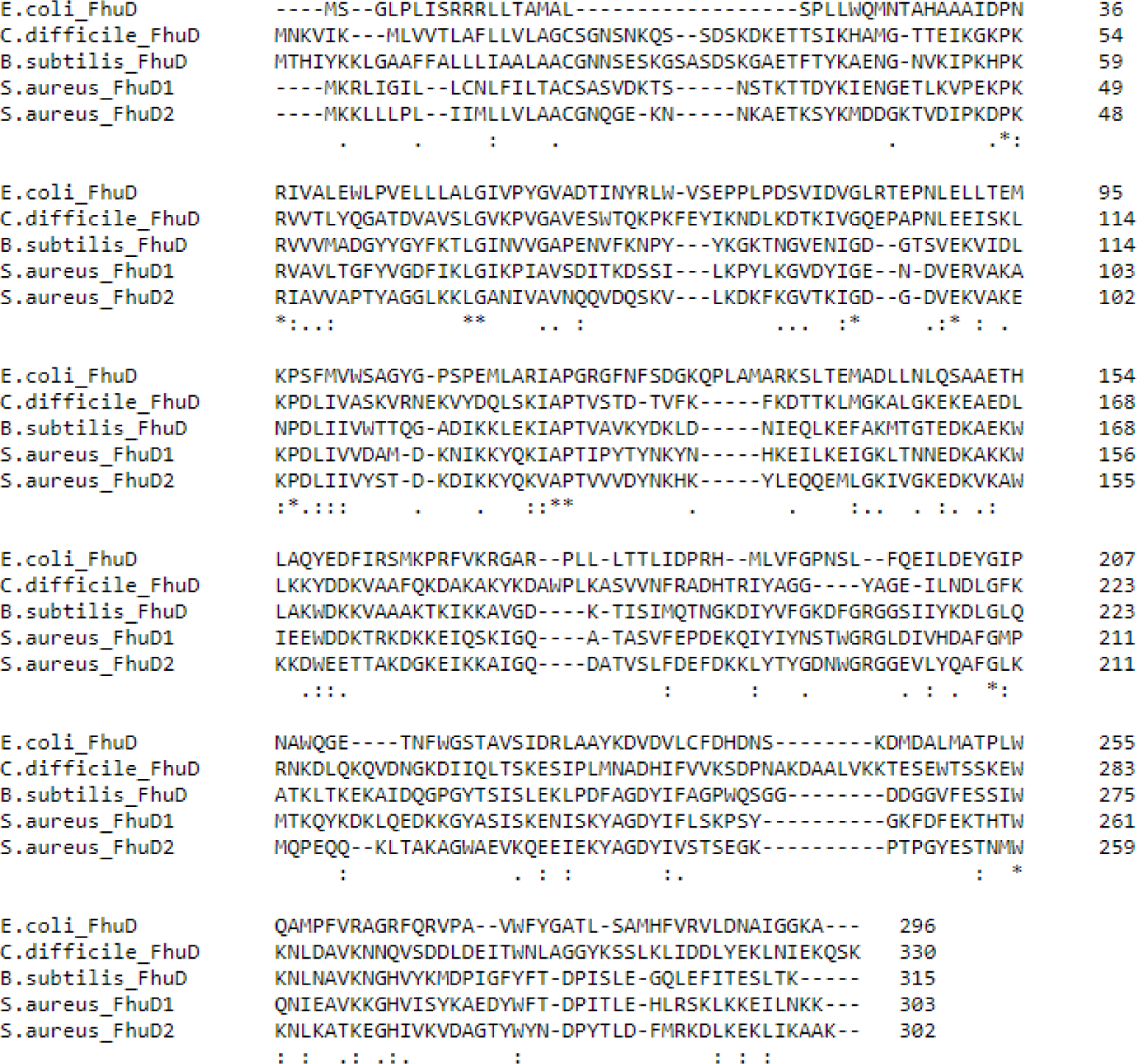
Clustal Omega alignment of FhuD.

**Supplemental 3.**
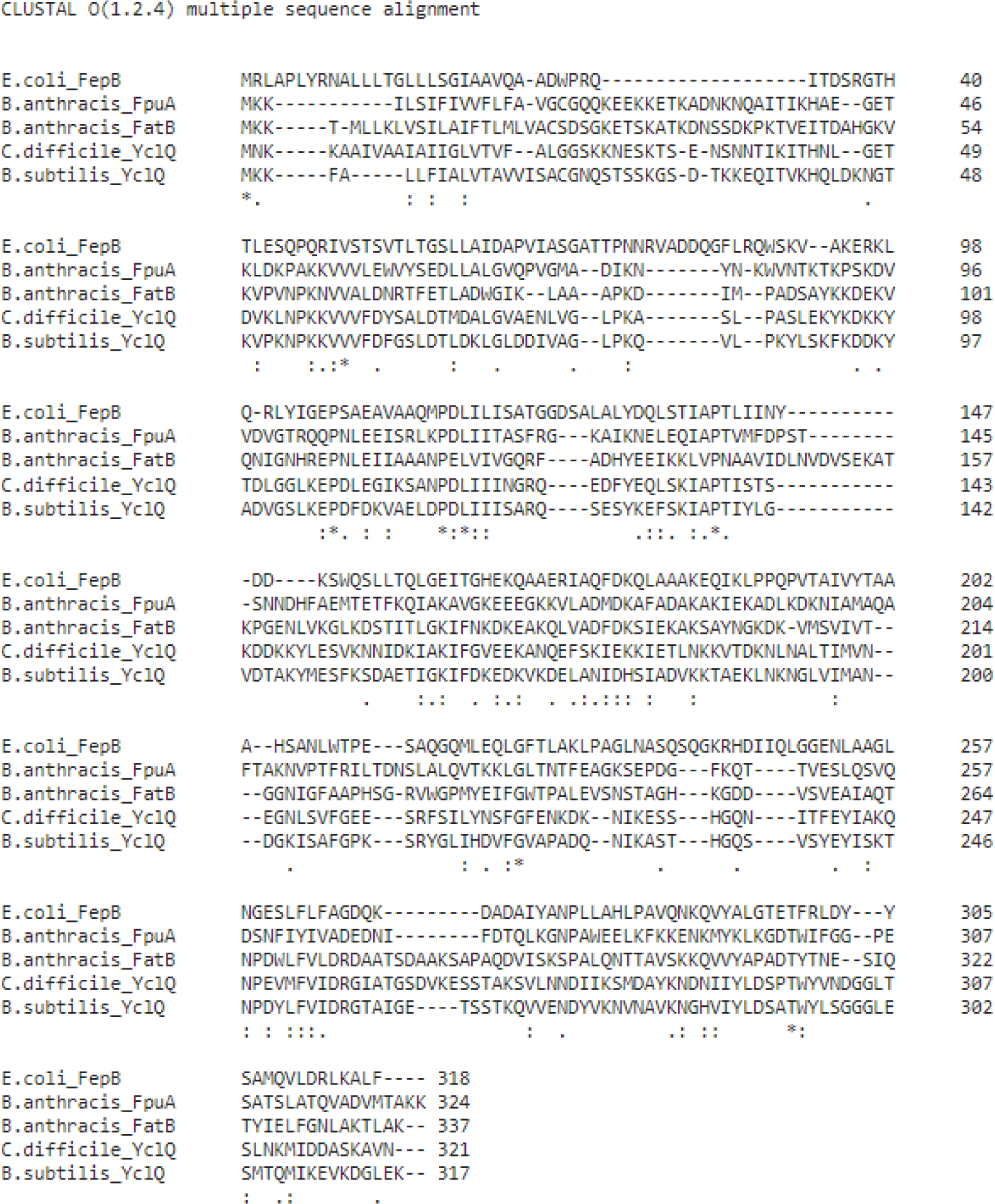
Clustal Omega alignment of YclQ.

**Supplemental 4.**
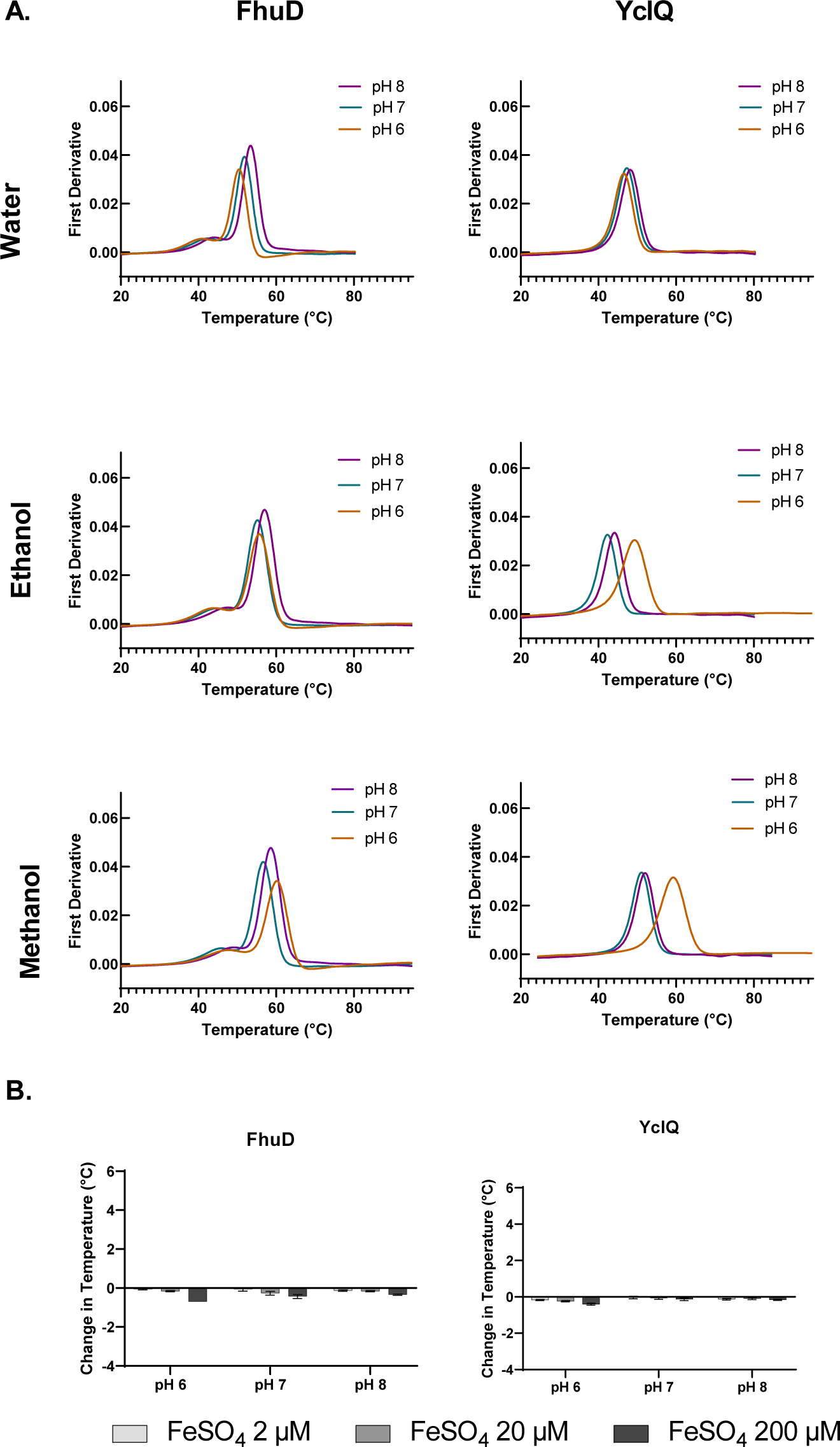
Controls from thermal shift experiments. Thermal unfolding of traces of siderophore binding protein (10 μM) across pH 6-8 in the medium used to resuspend the various siderophores (A). Purified FhuD and YclQ (10 μM) with increasing concentrations of iron (2-200 μM) (B).

**Supplemental 5.**
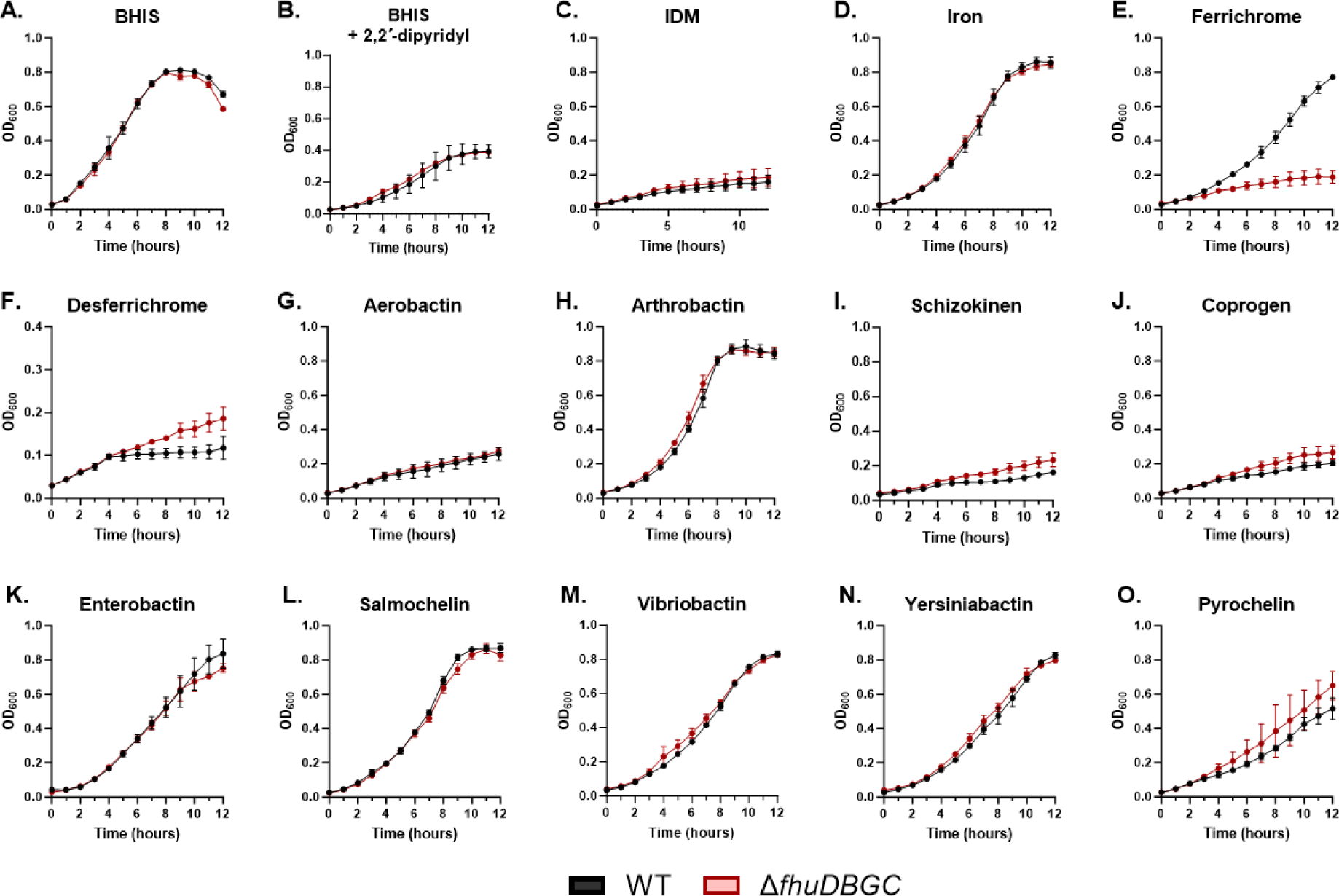
Reduced growth of *ΔfhuDBGC* is specific to ferrichrome (full growth curves). *C. difficile* 630 (black) and its isogenic *fhuDBGC* transporter mutant (red) was grown in iron starved conditions (BHIS + 75 μM 2’2’-dipyridyl) and adjusted to an OD 0.5 before diluting 1:10 into BHIS (A), BHIS supplemented with 75 μM 2’2’-dipyridyl (B), IDM (C), IDM supplemented with 2 μM FeSO_4_ (D) or IDM supplemented with 2 µM of siderophore pre-loaded with iron (E-O). Bacterial growth was assessed by determining the OD_600_ hourly for 12 hours. Data are presented as the mean ± standard deviation of three technical replicates. Data are representative of three biological replicates.

**Table S1.** Plasmids and Strains used in the study.

| Name | Description | Source or Reference |
| --- | --- | --- |
| **Plasmids** |  |  |
| pET24a+ | Overexpression plasmid | MilliporeSigma (Novagen) |
| pJH55 | pET24a Strep-Tactin tagged FhuD extracellular | This study |
| pMSR0 | toxin mediated allelic exchange | (12) |
| pJLH123 | pMSR0 with flanks targeting *fhuDBGC* (*Cd630_28780-28740*) | This study |
| pDSW1728 |  | (15) |
| pJLH150 | pDSW1728 + $P_{native}$-*fhuDBGC* | This study |
| ***E. coli*** |  |  |
| DH5α | used for cloning | NEB |
| HB101 | conjugation helper strain carrying pRK24 | (13) |
| BL21 | used for protein expression | NEB |
| ***C. difficile*** |  |  |
| 630 | Wild-type strain |  |
| 630 pJLH70 | 630 with CRISPRi targeting *fhuD* | This study |
| 630 Δ*fhuDBGC* | deletion of *fhuDBGC*+*MATE* | This study |
| 630 pDSW1728 | 630 with vector control | This study |
| 630 pJLH150 | 630 deletion of *fhuDBGC*+*MATE* with *fhuDBGC*+*MATE* complementation | This study |

**Table S2.**
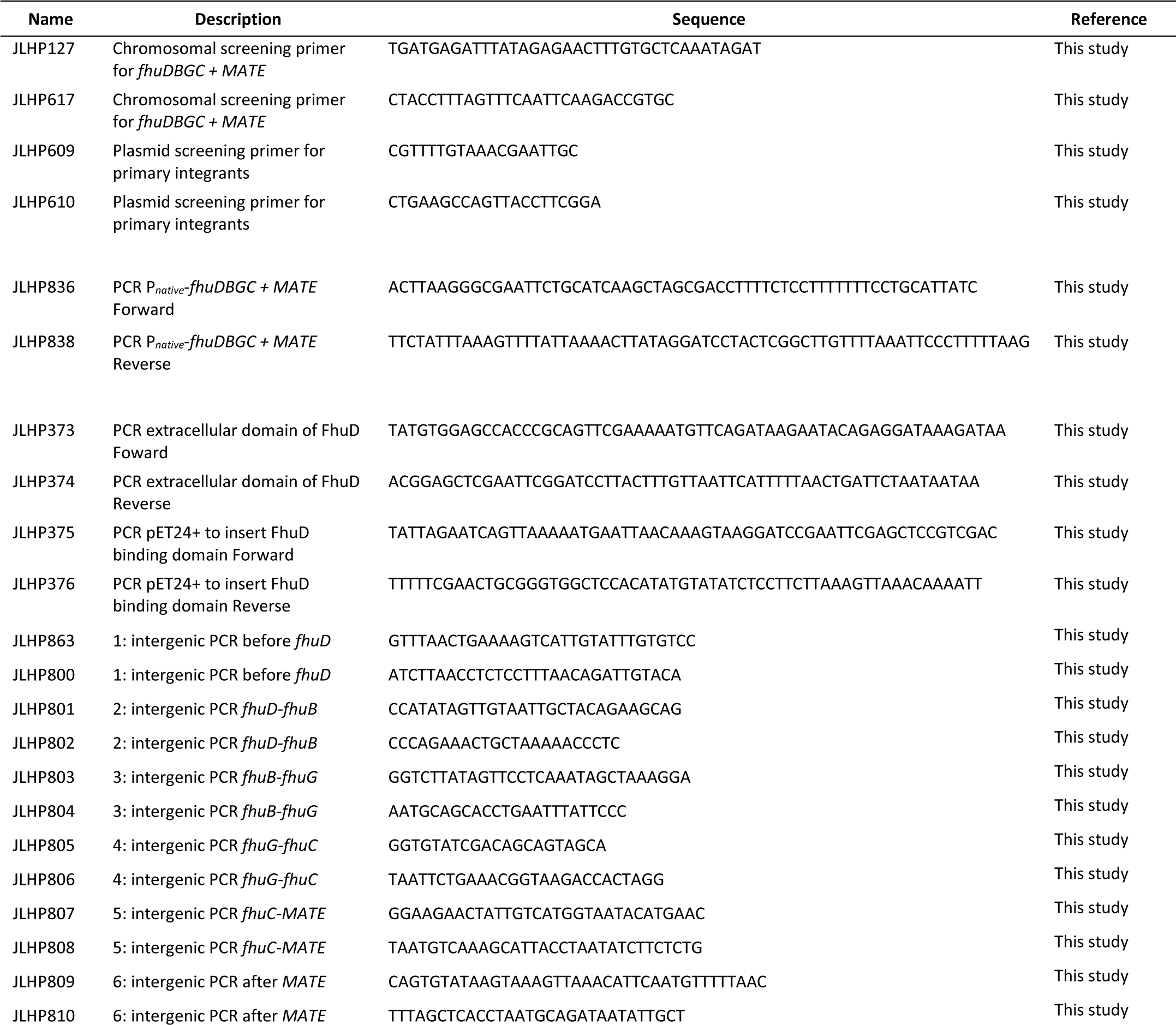
Primers used in the study.

**Table S3.**
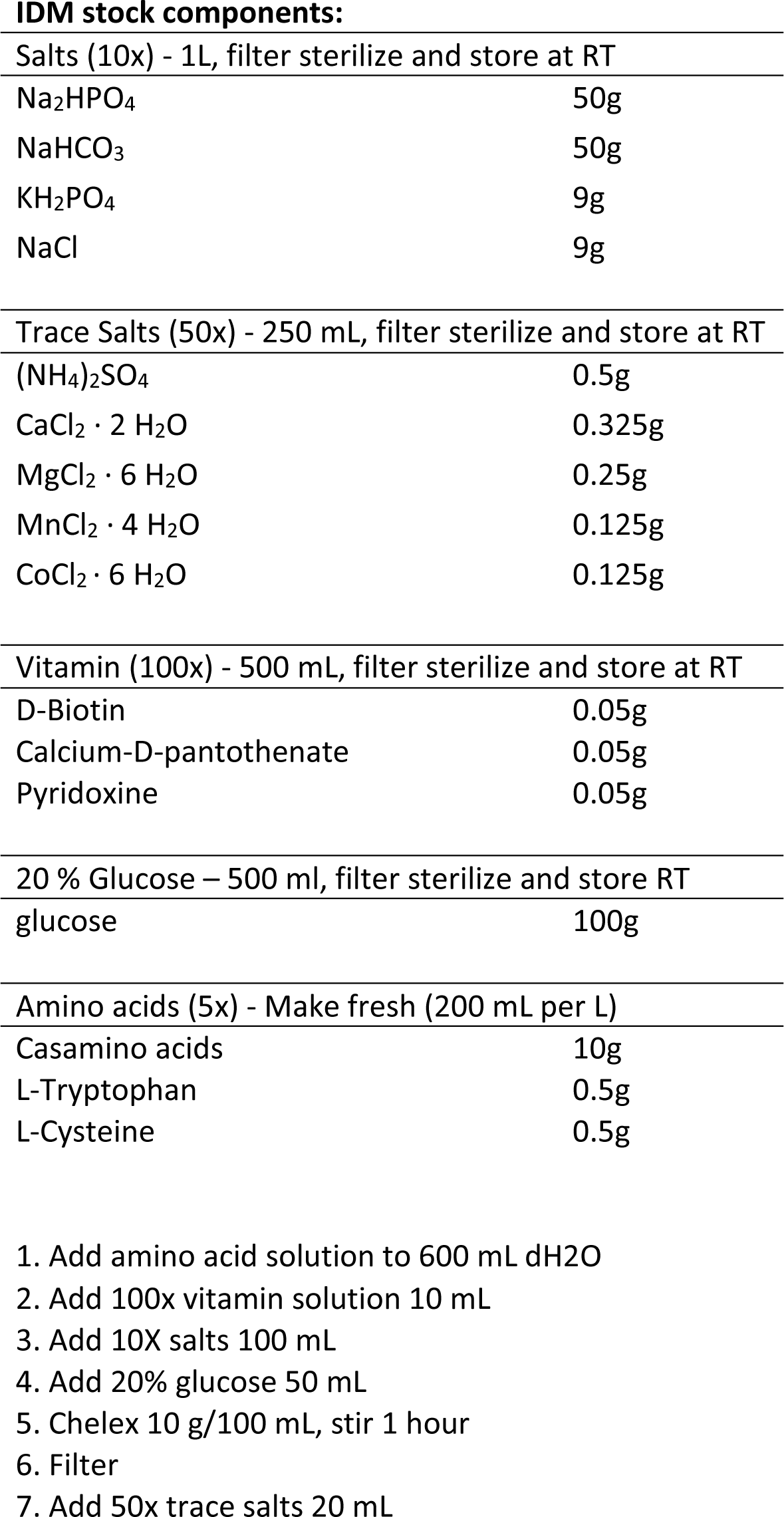
*C. difficile* Iron Depleted Media (IDM)

**IDM stock components:**
|  |  |
| --- | --- |
| Salts (10x) - 1L, filter sterilize and store at RT |  |
| Na<sub>2</sub>HPO<sub>4</sub> | 50g |
| NaHCO<sub>3</sub> | 50g |
| KH<sub>2</sub>PO<sub>4</sub> | 9g |
| NaCl | 9g |
| Trace Salts (50x) - 250 mL, filter sterilize and store at RT |  |
| (NH<sub>4</sub>)<sub>2</sub>SO<sub>4</sub> | 0.5g |
| CaCl<sub>2</sub> · 2 H<sub>2</sub>O | 0.325g |
| MgCl<sub>2</sub> · 6 H<sub>2</sub>O | 0.25g |
| MnCl<sub>2</sub> · 4 H<sub>2</sub>O | 0.125g |
| CoCl<sub>2</sub> · 6 H<sub>2</sub>O | 0.125g |
| Vitamin (100x) - 500 mL, filter sterilize and store at RT |  |
| D-Biotin | 0.05g |
| Calcium-D-pantothenate | 0.05g |
| Pyridoxine | 0.05g |
| 20 % Glucose – 500 ml, filter sterilize and store RT |  |
| glucose | 100g |
| Amino acids (5x) - Make fresh (200 mL per L) |  |
| Casamino acids | 10g |
| L-Tryptophan | 0.5g |
| L-Cysteine | 0.5g |

1. Add amino acid solution to 600 mL dH<sub>2</sub>O 2. Add 100x vitamin solution 10 mL 3. Add 10X salts 100 mL 4. Add 20% glucose 50 mL 5. Chelex 10 g/100 mL, stir 1 hour 6. Filter 7. Add 50x trace salts 20 mL

